# Leaf size determines damage- and herbivore-induced volatile emissions in maize

**DOI:** 10.1101/2024.11.14.623649

**Authors:** Jamie M. Waterman, Tristan M. Cofer, Ophélie M. Von Laue, Pierre Mateo, Lei Wang, Matthias Erb

## Abstract

Stress-induced plant volatiles play an important role in mediating ecological interactions between plants and their environment. The timing and location of the inflicted damage is known to influence the quality and quantity of induced volatile emissions. However, how leaf characteristics and herbivore feeding behavior interact to shape volatile emissions is not well understood. Using a high-throughput volatile profiling system with high temporal resolution, we examined how mechanical damage and herbivore feeding on different leaves shapes plant-level volatile emission patterns in maize. We then tested feeding patterns and resulting consequences on volatile emissions with two generalist herbivores (*Spodoptera exigua* and *Spodoptera littoralis*), and assessed whether feeding preferences are associated with enhanced herbivore performance. We found maize seedlings emit more volatiles when larger leaves are damaged. Larger leaves emitted more volatiles locally, which was the determining factor for higher plant-level emissions. Surprisingly, both *S. exigua* and *S. littoralis* preferentially consumed larger leaves, and thus maximize plant volatile emission without apparent growth benefits. Together, these findings provide an ecophysiological and behavioral mechanism for plant volatile emission patterns, with potentially important implications for volatile-mediated plant-environment interactions.

## Introduction

Time and space are important factors that shape plant defense expression and plant-environment interactions (Waterman *et al*., 2019; Hunziker *et al*., 2021). It is well known that discrete tissues, especially those with temporally separated development, exhibit specific defense capacities (Köhler *et al*., 2015; Maag *et al*., 2015; Maag *et al*., 2016; Hunziker *et al*., 2021). In general, evidence suggests that younger plant tissues are typically more well-defended (Ohnmeiss & Baldwin, 2000; Barton *et al*., 2010). While this is often true for inducible defenses, in some plants, such as maize, older tissues tend to have higher levels of constitutive chemical defenses (Köhler *et al*., 2015). As such, the specific defense, as well as the nature of its metabolism, are critical considerations. Additionally, spatiotemporal differences in inducible plant defense chemistry often depend on the specific mode of induction. In maize treated with ZmPep3, a peptide associated with regulating responses to wounding, older maize leaves respond more strongly, however, when exposed to herbivore-induced plant volatiles, younger leaves are more responsive (Wang *et al*., 2023). Even within the same leaf and over relatively short temporal time scales (hours), the timing and position of damage can strongly influence both qualitative and quantitative features of damage-induced responses (Mithöfer *et al*., 2005; Maag *et al*., 2016). Tissue-specific patterns may also be temporally dependent. Specifically, in cotton, young leaves from younger plants produced more terpenoids when mechanically damaged in comparison to young leaves in older plants (Eisenring *et al*., 2017). Similarly, damage to plants at different ontogenetic stages is also an important consideration. In lima bean, tolerance to herbivory through compensatory growth was observed in plants that were previously damaged (primed) during the seedling stages of development and not observed if plants were primed during juvenile and reproductive stages (Bustos-Segura *et al*., 2022).

An important damage-induced plant defense is the production of damage-induced volatiles. These volatiles are particularly versatile, as they are biosynthesized within tissues, where they can act as a direct defense against biotic stress (Maurya *et al*., 2020; Chen *et al*., 2021; Chen *et al*., 2023). These volatiles are also emitted into the environment, where they have broad ecological relevance (Escobar-Bravo *et al*., 2023). Plant-derived volatiles can serve as within-plant signaling molecules (Heil & Silva Bueno, 2007), attract herbivore natural enemies (Schnee *et al*., 2006; Turlings & Erb, 2018), influence herbivore oviposition (Hajdu *et al*., 2024), mediate plant-pollinator interactions (Groen *et al*., 2016) and mediate plant-plant interactions (Wang *et al*., 2023; Waterman *et al*., 2024).

Damage-induced volatiles can be manipulated by herbivores, which employ several strategies to reduce plant volatile emissions, potentially minimizing detection from natural enemies such as predators and parasitoids. For example, the saliva of the caterpillar *Helicoverpa zea*, which can be secreted onto plant tissues during feeding, contains the enzyme glucose oxidase, which causes stomatal closure and reduced the emission of some volatiles (Lin *et al*., 2021). Conversely, herbivores can also betray themselves by attracting natural enemies, for example *Manduca sexta* oral secretions decrease the *(Z)*/*(E)* ratio of green leaf volatiles emitted by plants during feeding, which increases herbivore attractiveness to predators (Allmann & Baldwin, 2010). While herbivore feeding patterns can influence plant phenotypes, within-plant spatiotemporal phenotypic variation can also influence herbivore feeding behavior, and in turn shape wider plant-environment interactions (Bellec *et al*., 2022; Orrock *et al*., 2022). While plant traits have been shown to influence herbivore behavior, broadly applicable patterns are difficult to identify and many herbivore-feeding patterns may be explained by processes and environmental features independent of the plant (The Herbivory Variability Network* † *et al*., 2023; Pan & Wetzel, 2024). Although volatile emissions have been shown to positively correlate with damage extent (Schmelz *et al*., 2003; Rodriguez-Saona *et al*., 2009), qualitative damage patterns are also known to shape volatile emission profiles (Mithöfer *et al*., 2005; Lange *et al*., 2020). Importantly, the degree to which herbivore feeding patterns and preferences align with inherent spatiotemporal differences in volatile induction is unclear.

The aim of our study was to test how damage done to different leaves affects the emission of ecologically relevant induced plant volatiles at the plant level, and to identify the mechanistic basis for these differences. Further, to better understand the potential implications of these differences, we investigated the feeding patterns of two lepidopteran herbivore species, and the resulting impact on herbivore-induced volatile emissions. We tested a number of biochemical, chemical and morphological leaf traits across developmental stages, and manipulated leaf size to build a mechanistic understanding of what drives differences in emission between leaves. Additionally, we determined that herbivores preferentially fed on larger leaves without any apparent performance benefit, and thus maximized plant volatile emissions. Together, these findings reveal leaf size as an important determinant of plant-level volatile emissions and demonstrate that two generalist herbivores feed in a way that maximizes rather than minimizes leaf volatile emissions, without apparent direct benefits.

## Materials and Methods

### Plant and insect growth

V2- and V3-stage maize seedlings (*Zea mays*, B73) were used throughout this study. At the V2 stage, maize has four leaves: two are fully developed, one is expanding and one is emerging. At the V3 stage maize has 5 leaves: three are fully developed, one is expanding and one is emerging. At these growth stages, maize plants are often attacked by herbivores such as *Spodoptera* spp. (van den Berg *et al*., 2021). Maize plants were grown in commercial potting soil (Selmaterra, BiglerSamen, Switzerland) in 180 ml pots. Plants were grown in greenhouse conditions and supplemented with artificial lights (*ca*. 300 µmol m^−2^ s^−1^). The greenhouse was kept at 22 ± 2 °C, 40–60% relative humidity, with a 14 h : 10 h, light : dark cycle. *Spodoptera exigua* (Frontier Agricultural Sciences, USA) and *Spodoptera littoralis* (University of Neuchatel, Switzerland) were reared from eggs on artificial diet (Maag *et al*., 2014) and used for experiments when they reached the 4^th^ instar stage. All experiments were conducted between the hours of 10:00 and 20:00 under full-light conditions.

### Plant damage

To mimic dynamic patterns, we damaged plants three times in *ca.* 30 min intervals with a hemostat. Each wound was *ca.* 10 mm^2^. Each damage event consisted of two wounds, one on each side of the midvein. We chose damage treatments without application of larval oral secretions to obtain generalizable induction patterns without species-specific induction and suppression of volatiles (Lange *et al*., 2020). The first damage event was at the base of the leaf, the second in the middle, and the third at the tip. The extent and relative spacing of damage was identical across leaves, so that for each leaf the tip, middle and base of each leaf was damaged. Damage was either inflicted on leaves of intact plants or detached leaves as specified in the text and figure legends.

### Detached leaf assay

Leaves were dissected following the protocol of Wang et al., 2023. In brief, leaves were cut at the base from maize seedlings while submerged in Milli-Q water. The cut base of leaves remained submerged in water for 2 hr to allow for tissues to recover from dissections, prior to damage treatments. Volatile emissions from individual leaves were measured the same way as intact seedlings.

### Volatile sampling

Entire seedlings or detached leaves were placed in transparent glass chambers (Ø×H 12 × 45 cm) that were sealed other than an clean airflow inlet and an outlet. Clean air was supplied at a flow-rate of 0.8 L min^−1^. Volatiles were measured with a high-throughput platform comprising of a proton transfer reaction time-of-flight mass spectrometer (PTR-ToF-MS; Tofwerk, Switzerland) and a custom-made automated headspace sampling system (Gonin *et al*., 2018). The outlet of the chamber was accessible to the autosampler/PTR-ToF-MS system. The PTR-ToF-MS system drew air at 0.1 L min^−1^. Between samples, a zero-gas measurement was performed for 3 s to flush the system. At each time point (as indicated by the x-axis of the respective figure), volatiles were continuously measured for 25-30 s and averaged to a single mean per sample. When total emission over the experimental period is reported, the frequency of each measurement is specified in the respective figure legend. Complete mass spectra (0-500 m/z) were recorded in positive mode at *ca.* 10 Hz. The PTR was operated at 100 °C and an E/N of approximately 120 Td. The volatile data extraction and processing were conducted using Tofware software package v3.2.2 (Tofwerk, Switzerland). Protonated compounds were identified based on their molecular weight + 1. During volatile collection, LED lights (DYNA, heliospectra) were placed *ca.* 80 cm above the glass cylinders and provided light at 300 μmol m^−2^ s ^−1^. Identical light : dark cycle timing as in the greenhouse for plant growth was used. To calculate ‘total’ volatile emissions, we took the integral of the emission curve over time, in other words, the sum of each individual volatile measurement time point over the experimental period.

### Gene expression

1.5 hr after the first leaf damage, each leaf (leaf 2, 3 and 4) of each plant was harvested and flash frozen on liquid nitrogen. Total RNA extraction and purification, genomic DNA removal, cDNA synthesis and quantification of gene expression were conducted identically to Waterman et al., 2024. Quantitative reverse transcription polymerase chain reaction (qRT-PCR) was performed using ORA SEE qPCR Mix (highQu GmbH, Germany) on an Applied Biosystems QuantStudio 5 Real-Time PCR system. The normalized expression (NE) values were calculated as in Waterman et al., 2024 using ubiquitin (UBI1) as the reference gene. Gene identifiers and primer sequences are listed in Supplementary Table 1.

### HDMBOA-Glc quantification

24 hr after the first leaf damage, each leaf (leaf 2, 3 and 4) of each plant was harvested and flash frozen on liquid nitrogen. Approximately 100 mg of fresh frozen, finely ground leaf was extracted in 1 mL acidified MeOH/H_2_O 70:30 v/v spiked with 0.1% formic acid. Extracts were vortexed for 10 seconds and centrifuged at 13000 rpm for 20 min at 10 °C. The supernatant was removed and used for subsequent quantification. HDMBOA-Glc profiling was performed with an Acquity i-Class UPLC system equipped with a photodiode array detector (PDA) and a single quadrupole detector (QDa) with electrospray source (Waters, USA). Gradient elution was performed on an Acquity BEH C18 column (1.7 μm, 2.1 × 100 mm, Waters, USA).The elution conditions were as follows: 90-70% A over 3 min, 70-60% A over 1 min, 40-100% B over 1 min, holding at 100% B for 2.5 min, holding at 90% A for 1.5 min where A = 0.1% formic acid/water and B = 0.1% formic acid/acetonitrile. The flow rate was 0.4 mL/min. The temperature of the column was maintained at 40 °C, and the injection volume was 1 μL. Absolute quantities were determined using standard curves of the corresponding pure HDMBOA-Glc.

### Stomatal density

Clear nail polish was painted onto either the base, middle or tip of the abaxial and adaxial side of each leaf. Once hardened, the polish was removed with double-sided sticky tape, creating an epidermal imprint. Imprints were mounted onto glass microscope slides and imaged with a Leica DM2500 optical microscope coupled to an HD digital microscope camera (Leica MC170 HD; Leica Microsystems, Germany). The density of stomata on each imprint was calculated based on the average of three distinct and non-overlapping fields of view.

### Herbivore feeding on intact plants

To assess damage patterns on intact V2-stage maize seedlings, a single *S. exigua* or *S. littoralis* larva was placed on either leaf 2 or leaf 3. Herbivores were allowed to feed for 6 hr. At the end of the feeding period, damaged area (percentage of total leaf) of each leaf was estimated visually. To convert this estimated value on a per-leaf basis into the percentage of total plant damage, a subset of plants was kept undamaged. Using ImageJ (National Institutes of Health, USA), the average surface area of each leaf on these control plants was used as a reference, and the percent area contributed by each individual leaf to the total surface area of the plant was calculated. In other words, the estimated percentage of a given leaf removed by herbivores was multiplied the percentage of the total area of all leaves made up by said leaf.

### Herbivore feeding and performance on individual leaves

To assess herbivore feeding on detached leaf 2 and leaf 3, leaves were dissected as previously described (see *detached leaf assay*). One 4^th^ instar larva was placed at the base of each leaf and allowed to feed for 6 hr. The area of tissue damage was calculated using ImageJ and methods described in (Waterman *et al*., 2021). To calculate the biomass of tissue consumed we multiplied the area of damage by the leaf mass per area (LMA), which is descried as the ratio of mass per unit area (KATTGE *et al*., 2011). This was calculated for leaf 2 (LMA = 0.0031, *n* = 6) and leaf 3 (LMA = 0.0025, *n* = 6). To assess the relative growth rate of herbivores on detached leaves, larvae were placed on leaves in an identical fashion to described above, however were allowed to feed for 48 hr, with leaves being replaced at the 24 hr mark to prevent desiccation. Relative growth rate was calculated as: ((final larval weight – initial larval weight) / initial larval weight) / # of days of feeding, as previously described (Waterman *et al*., 2021). 48 hr feeding periods were chosen to approximate the time it would take a 4^th^ instar *Spodoptera* larva to consume leaf 2 or leaf 3, while also allowing robust inferences regarding the quality of plant material for larval development (Himanen *et al*., 2009; Paul-Victor *et al*., 2010; Underwood, 2012; Lin *et al*., 2020; Waterman *et al*., 2021).

### Statistical analyses

All statistical analyses were conducted in R version 4.2.2 (R Core Team, 2022). Gene expression and stomatal density were analyzed using two-way ANOVA (Type = II). Gene expression was analyzed on log-transformed normalized expression values. HDMBOA-Glc was analyzed with one-way ANOVA within each leaf for plants where leaf 2 and leaf 3 were damaged. As data from plants with leaf 4 damage did not meet the assumptions of ANOVA, differences in HDMBOA-Glc between leaves of these plants were determined using a Kruskal-Wallis test. Differences in detached leaf biomass were also analyzed with one-way ANOVA. Where necessary, to obtain heteroscedasticity-consistent standard errors, White-adjusted ANOVAs were used (White, 1980). To determine leaf-damage area by herbivores in intact seedlings, as the assumptions of ANOVA were not met, Kruskal-Wallis tests were used. For herbivore feeding and performance assays in detached leaves, Welch’s two-sample t-tests were used to compare leaf 2 and leaf 3. Statistical test summaries are included as supplemental information (Supplemental Tables 2 and 3).

## Results

### Plant-level volatile emissions are determined by damage position

In order to determine how damage position impacts induced plant-level volatile emissions, we damaged each leaf of a V2-stage maize seedling individually (one damaged leaf per plant; Fig 1A) and measured induced volatile emissions. Green leaf volatile emissions, which are strongly localized to wound sites (Matsui, 2006), were comparable across treatments, with the exception of plants where the youngest leaf (leaf 4) was damaged, which resulted in, generally, lower green leaf volatile emissions (Fig 1B). Plant-level emissions of terpenes and indole were strongly influenced by damage location. Plants damaged on leaf 3 had the highest emission levels of monoterpenes [C_10_H_17_^+^, m/z = 137.13], sesquiterpenes [C_15_H_25_^+^, m/z = 205.20], and 4,8,12-trimethyltrideca-1,3,7,11-tetraene (TMTT) [C_16_H_27_^+^, m/z = 219.21], 4,8-dimethylnona-1,3,7-triene (DMNT) [C_11_H_19_^+^, m/z = 151.15] and indole [C_8_H_8_N^+^, m/z = 118.07], followed by leaf 2 and leaf 4 (Fig 1C). Thus, which leaf is damaged is a strong determinant of plant-level volatile emissions.

**Figure 1.**
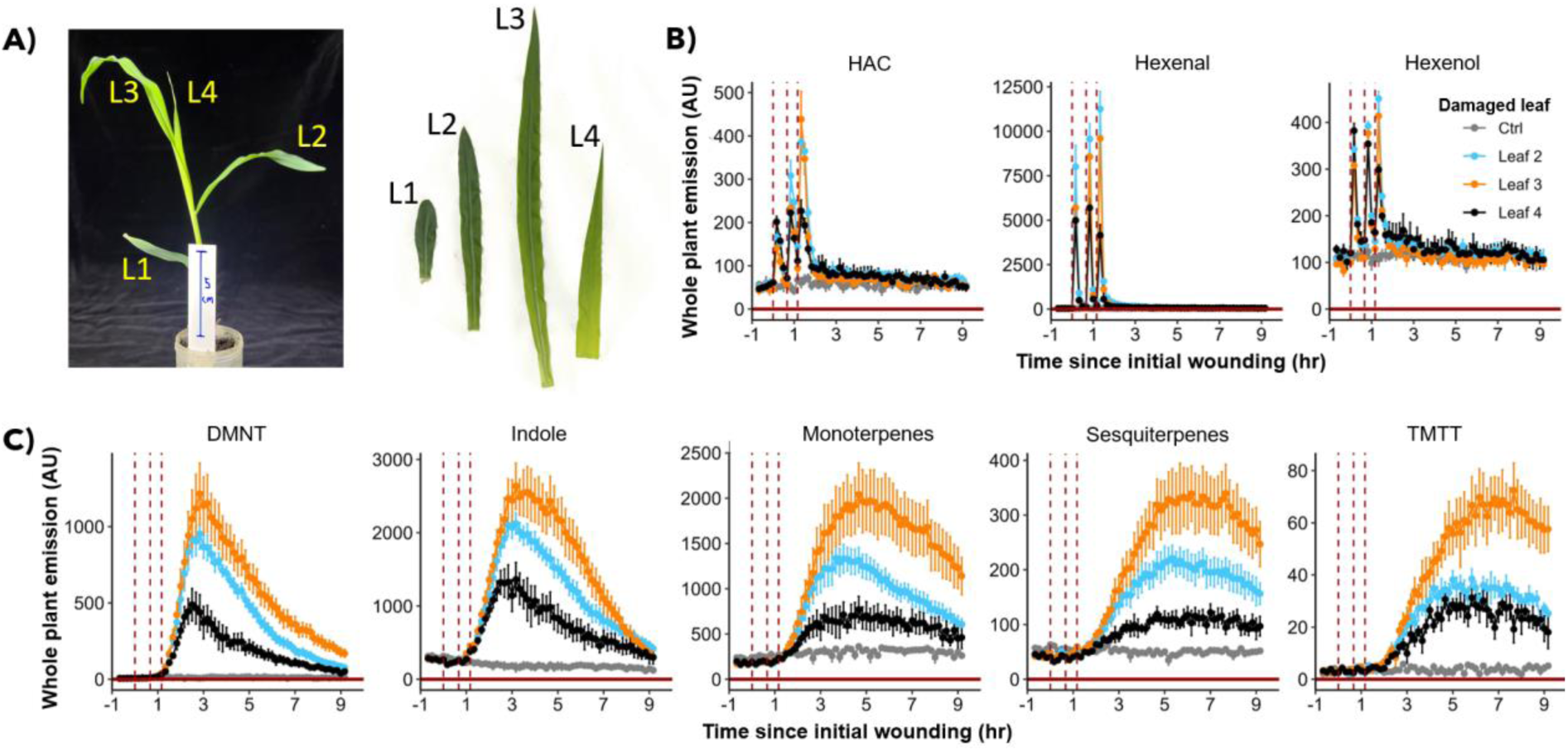
Plant-level volatile emissions depend on which leaves are damaged. A) The different leaves and respective sizes of intact V2-stage maize seedlings. Emission kinetics of B) green leaf volatiles and C) terpenes and indole are shown. Perforated vertical lines depict the timing of each mechanical damage entry. Abbreviations: HAC, Hexenyl acetate; DMNT, 4,8-dimethylnona-1,3,7-triene; TMTT, 4,8,12-trimethyltrideca-1,3,7,11-tetraene. Compounds were identified based on their molecular weight + 1, as all compounds were protonated. HAC: m/z = 143.11, Hexenal: m/z = 99.08, Hexenol: m/z = 101.0, DMNT: m/z = 151.15, Indole: m/z = 118.07, Monoterpenes: m/z = 137.13, Sesquiterpenes: m/z = 205.20, TMTT: m/z = 219.21. Colored points = mean ± SE. *n* = 5.

### Minimal differences in local damage inducibility of volatile biosynthesis genes and direct defenses

To test whether damage to the different leaves leads to differential local and systemic defense induction, which may determine leaf- and plant-level volatile emissions, we measured terpene and indole biosynthesis gene expression. Terpene synthase 2 and 10 (TPS2 and TPS10), as well as dimethylnonatriene/trimethyletetradecatetraene synthase (CYP92C5) are rate limiting for terpene production, and indole-3-glycerol phosphate lyase (IGL) is rate limiting for volatile indole biosynthesis (Frey *et al*., 2000; Richter *et al*., 2016; Block *et al*., 2019). We also measured HDMBOA-Glc levels as a major marker of non-volatile defense induction in maize to test whether spatial defense patterns may be generalizable (Li *et al*., 2018). Following damage to individual leaves, all tested genes were significantly induced both locally and systemically (Fig 2). Local induction was generally highest in leaf 2 and leaf 3 and lower in leaf 4 (Fig 2). Systemic induction was similar across treatments. HDMBOA-Glc was induced locally, but not systemically across all leaves at comparable concentrations, although the effect was only significant for leaf 2 and leaf 3 (Supplemental Fig 1). Thus, the differences in plant volatile emissions upon damage of cannot be explained by differential local and systemic inducibility.

**Supplemental Figure 1.**
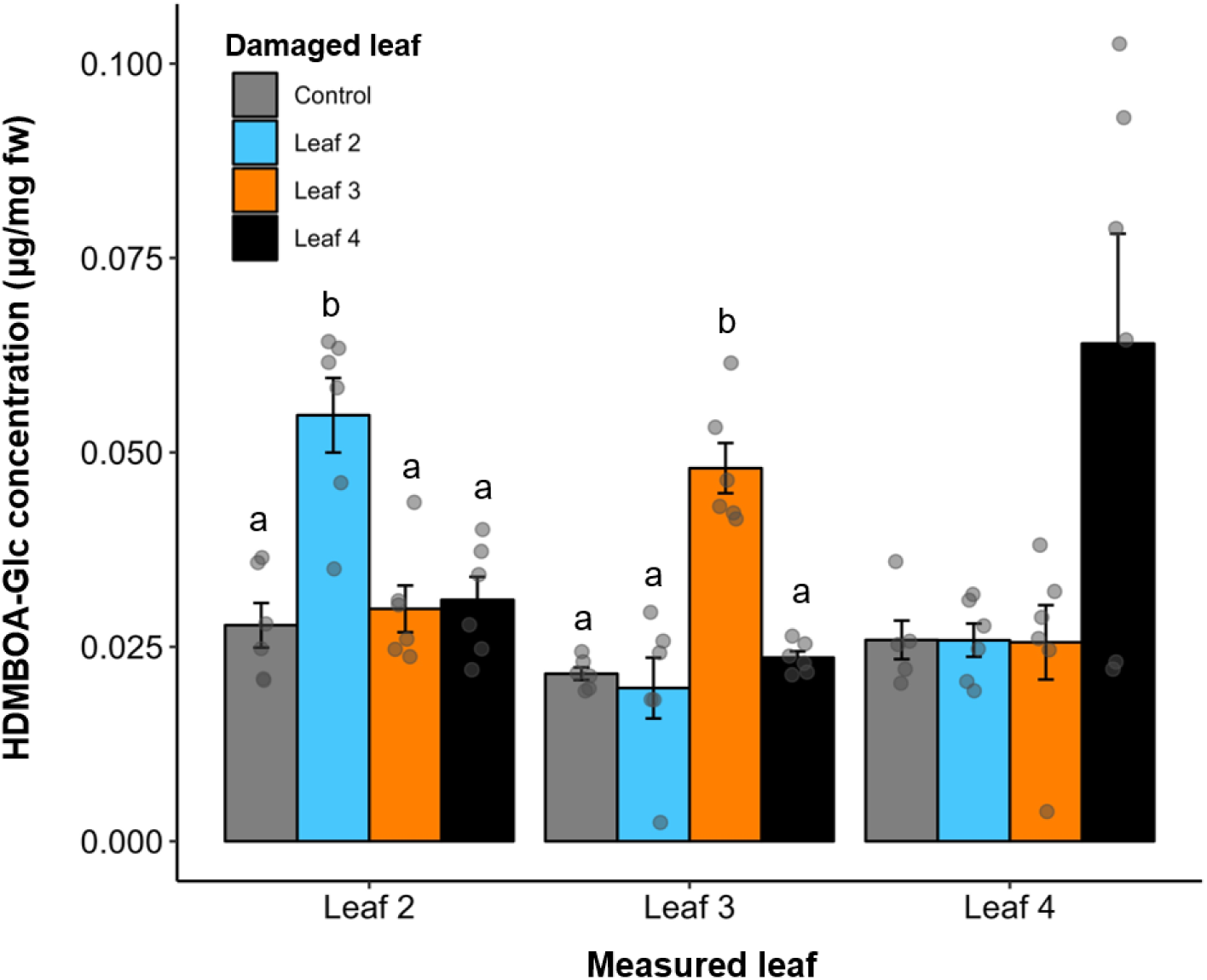
HDMBOA-Glc is similarly induced across leaves in intact V2-stage maize seedlings. Lowercase letters indicate significant differences between damage sites within a given leaf based on multiple comparisons tests. Grey points represent biological replicates. Bars = mean ± SE. *n* = 5-6.

**Figure 2.**
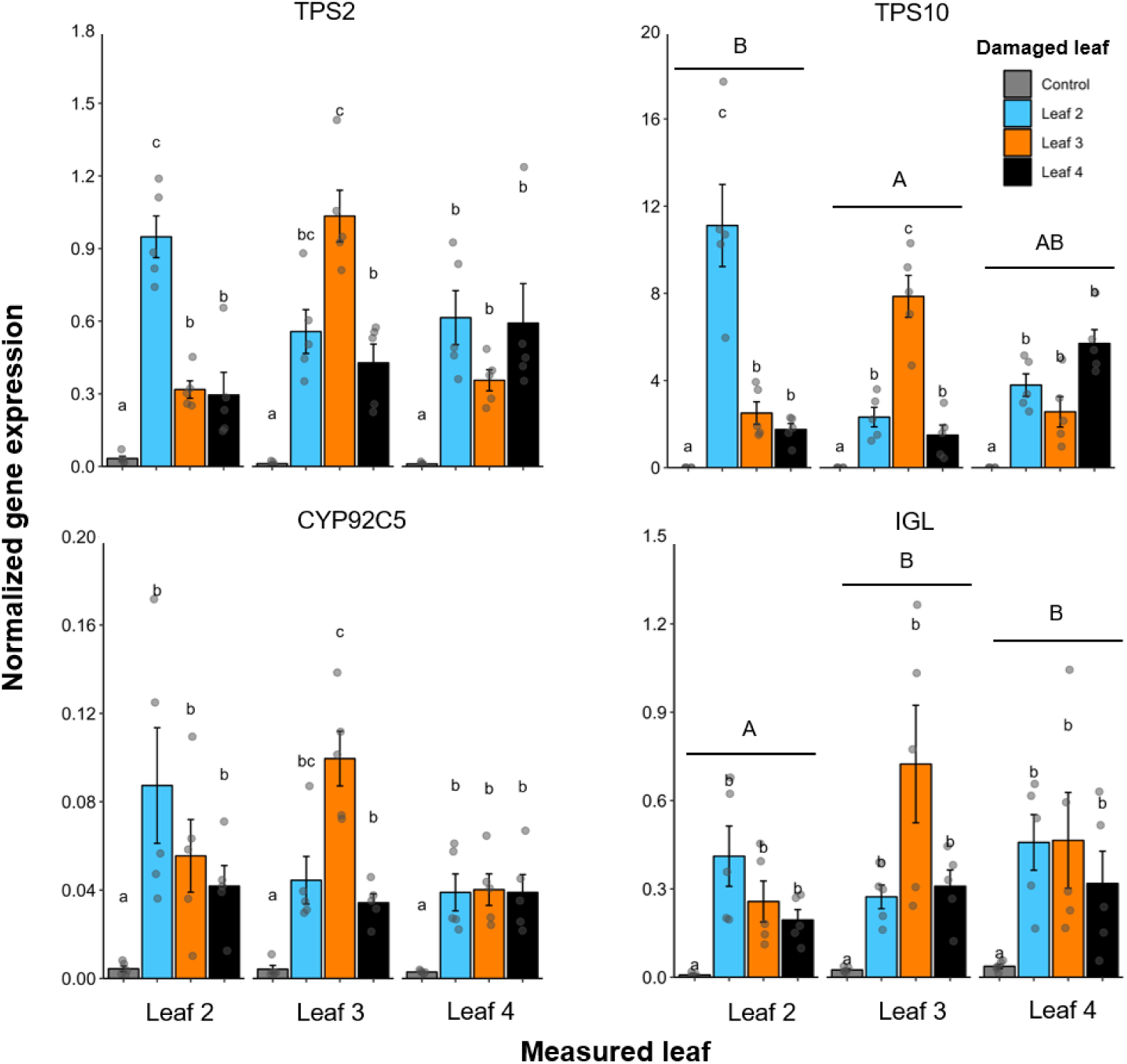
Minimal differences in volatile biosynthesis inducibility of different leaves in intact plants. Expression of terpene and indole biosynthesis genes are shown. Abbreviations: TPS2, terpene synthase 2; TPS10, terpene synthase 10; CYP92C5, Dimethylnonatriene/trimethyltetradecatetraene synthase, IGL, indole-3-glycerol phosphate lyase. Lowercase letters indicate significant differences between damage sites within a given leaf and uppercase letters indicate overall differences between leaves, regardless of which left was damaged (based on multiple comparisons tests). Grey points represent biological replicates. Bars = mean ± SE. *n* = 5.

### Plant volatile emissions strongly correlate with the size of the damaged leaf

One apparent difference between the leaves is their size, both in terms of biomass and surface area (Fig 1A). Thus, we hypothesized that larger leaves may result in higher volatile release upon damage. We used biomass as the principal metric of size, which is also highly correlated with leaf surface area across all leaves, and this correlation is maintained across developmental stages (Supplemental Fig 2). We found strong and significant correlations between the biomass of the damaged leaf and plant-level volatile emissions (Fig 3A). We also found similar correlations for leaf-level emissions in individual detached damaged leaves of V2- and V3-stage seedlings (Fig 3B; Supplemental Fig 3). There was also a significant linear relationship between volatile emissions and the surface area of the damaged leaf in both seedling stages (albeit slightly weaker than the relationship with damaged leaf mass; Supplemental Fig 4). Immature leaves (leaf 3 and 4 for V2-stage seedlings and leaf 4 and 5 for V3-stage seedlings) showed the strongest relationship between emission and size across the different leaves (Fig 3; Supplemental Fig 4). Interestingly, leaf 2 consistently showed higher volatile emissions than expected for its size in V2-stage seedlings, as did leaf 3 in V3-stage seedlings (Fig 3).

**Supplemental Figure 2.**
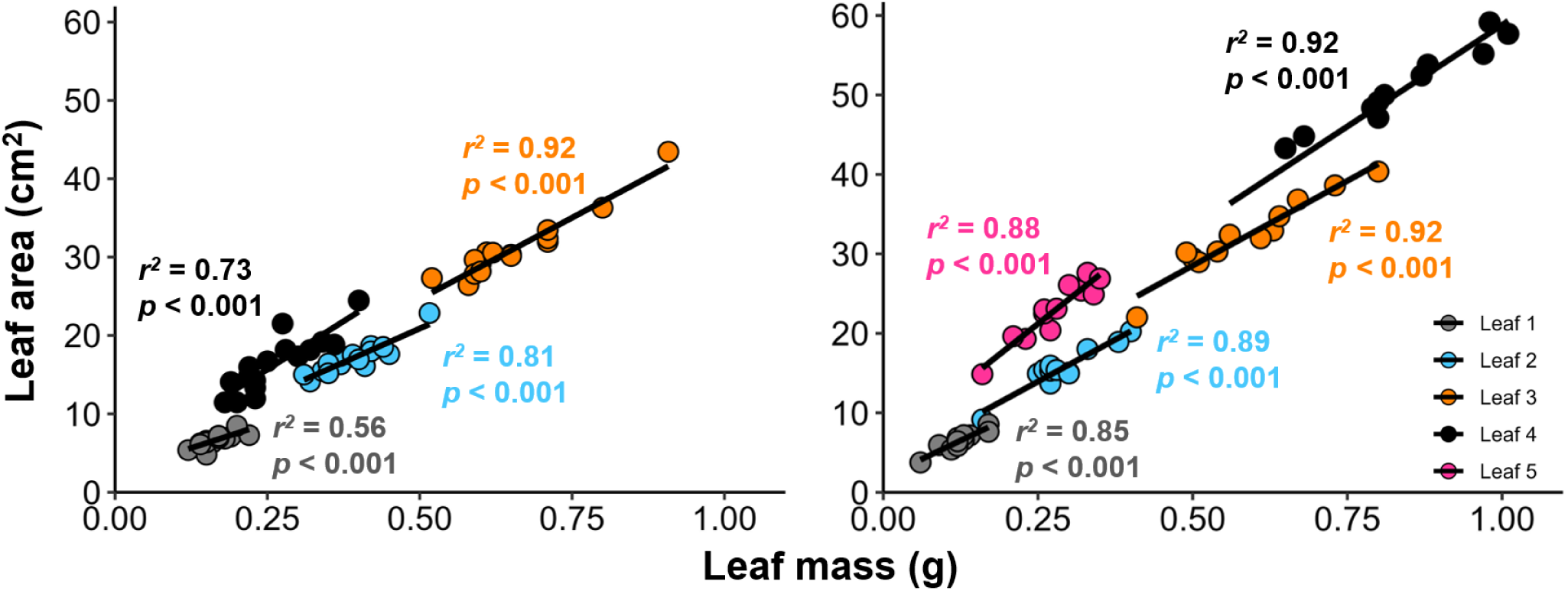
Strong positive relationship between leaf mass and leaf area in both V2-stage and V3-stage maize seedlings. Black lines represent linear regression for significant linear relationships for each leaf. Colored points depict biological replicates of different leaves. Plants used were undamaged. *n* = 12-16.

**Figure 3.**
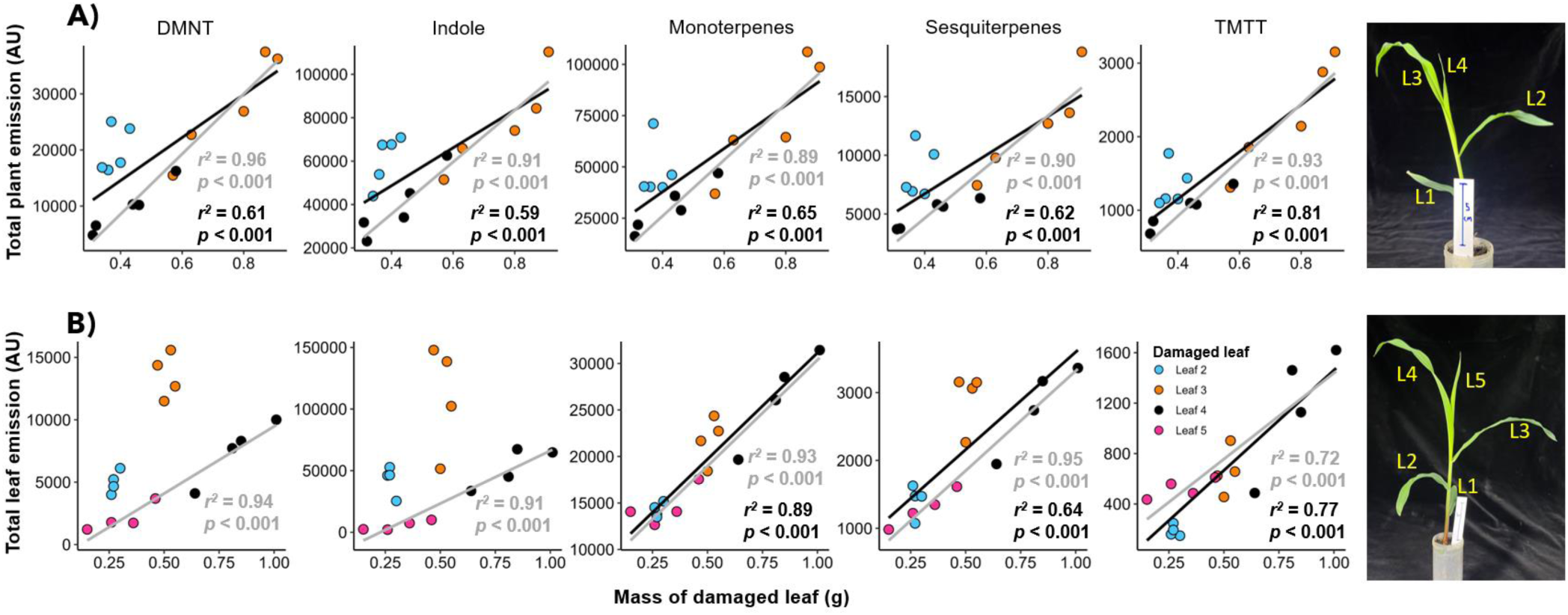
Whole-plant- and leaf-level volatile emissions strongly correlate with the mass of the damaged leaf. Colored points represent biological replicates and depict the relationship between total volatile emission (sum of individual measurement time points over 9 hr; see Materials and Methods, *Volatile sampling*) and leaf biomass in A) intact V2-stage seedlings and B) detached leaves of V3-stage seedlings. Individual measurements were taken every *ca.* 10 or 20 min for V2 and V3, respectively. Black lines represent linear regression for significant linear relationships between all leaves. Grey lines represent linear regression for significant linear relationships between immature leaves (leaf 3 and leaf 4 for V2-stage; leaf 4 and leaf 5 for V3-stage). Abbreviations: DMNT, 4,8-dimethylnona-1,3,7-triene; TMTT, 4,8,12-trimethyltrideca-1,3,7,11-tetraene. Compounds were identified based on their molecular weight + 1, as all compounds were protonated. DMNT: m/z = 151.15, Indole: m/z = 118.07, Monoterpenes: m/z = 137.13, Sesquiterpenes: m/z = 205.20, TMTT: m/z = 219.21. *n* = 5 for V2-stage seedlings, *n* = 4 for V3-stage leaves.

**Supplemental Figure 3.**
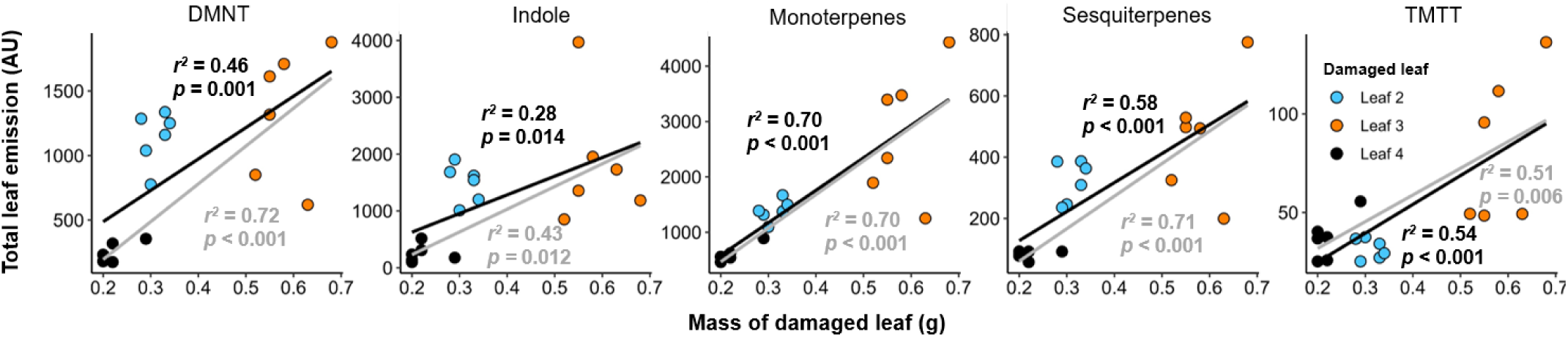
Detached maize leaves from V2-stage seedlings maintain the same emission-leaf size relationships as when leaves of intact plants are damaged. Colored points represent biological replicates and depict the relationship between total volatile emission (sum of individual measurement time points over 9 hr; see Materials and Methods, *Volatile sampling*) and leaf biomass. Individual measurements were taken every *ca.* 10 min. Black lines represent linear regression for significant correlations between all leaves. Grey lines represent linear regression for significant linear relationships between immature leaves (leaf 3 and leaf 4). Abbreviations: DMNT, 4,8-dimethylnona-1,3,7-triene; TMTT, 4,8,12-trimethyltrideca-1,3,7,11-tetraene. Compounds were identified based on their molecular weight + 1, as all compounds were protonated. DMNT: m/z = 151.15, Indole: m/z = 118.07, Monoterpenes: m/z = 137.13, Sesquiterpenes: m/z = 205.20, TMTT: m/z = 219.21. Error bars = SE. *n* = 6.

**Supplemental Figure 4.**
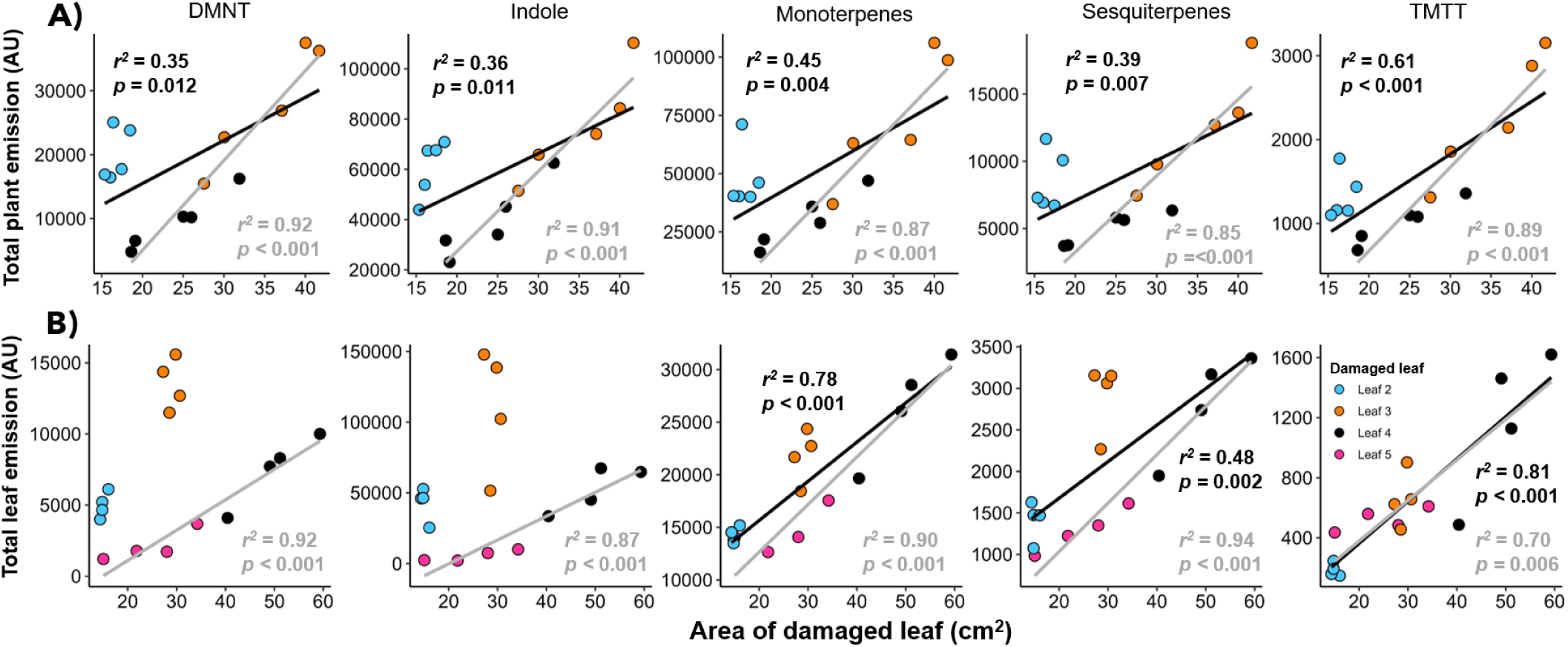
Relationships are maintained between volatile emissions and the area of damaged leaves across developmental stages. Colored points represent biological replicates and depict the relationship between total emission (sum of individual measurement time points over 9 hr; see Materials and Methods, *Volatile sampling*) of plant volatiles and leaf surface area in A) intact V2-stage seedlings and B) detached leaves of V3-stage seedlings. Individual measurements were taken every *ca.* 10 or 20 min for V2 and V3, respectively. Black lines represent linear regression for significant correlations between all leaves. Grey lines represent linear regression for significant linear relationships between immature leaves (leaf 3 and leaf 4 for V2-stage; leaf 4 and leaf 5 for V3-stage). Abbreviations: DMNT, 4,8-dimethylnona-1,3,7-triene; TMTT, 4,8,12-trimethyltrideca-1,3,7,11-tetraene. Compounds were identified based on their molecular weight + 1, as all compounds were protonated. DMNT: m/z = 151.15, Indole: m/z = 118.07, Monoterpenes: m/z = 137.13, Sesquiterpenes: m/z = 205.20, TMTT: m/z = 219.21. Error bars = SE. *n* = 5 for V2-stage seedlings, *n* = 4 for V3-stage leaves.

### Standardizing leaf area results in similar induced volatile emissions

In order to investigate whether leaf size is the driving factor for induced volatile emission, we compared entire detached leaves with leaves that were all cut to the biomass of leaf 4 (Fig 4A-C). Entire detached leaves showed similar emission kinetics to entire plants, where leaf 3 emitted the most volatiles (Fig 4D). When cut to equal size, emissions of damage-induced volatiles were similar between leaf 2 and 3 (Fig 4E). For all compounds except TMTT, both leaf 2 and leaf 3 emitted more volatiles than leaf 4 when cut to equal biomass. As terpene and indole emission are controlled by stomata, we measured stomatal densities between leaf 2 and leaf 3 and found no differences (Supplemental Fig 5). Maize-leaf development occurs on a gradient from the tip to base, with the tip being most developed and the base being least developed (Raissig *et al*., 2017; Wang *et al*., 2019). Accordingly, the base of leaves has fewer developed stomata than the other sections (Supplemental Fig 5). When cutting leaves to similar sizes, we only removed the bottom portion (with fewer on stomata for both leaves). There were no differences stomatal density between the middle and tip in leaves 2 and 3, and no differences in stomatal density were observed between leaf 2 and leaf 3 (Supplemental Fig 5), thus ruling out stomatal density as a driver of differential volatile emissions between these two leaves.

**Figure 4.**
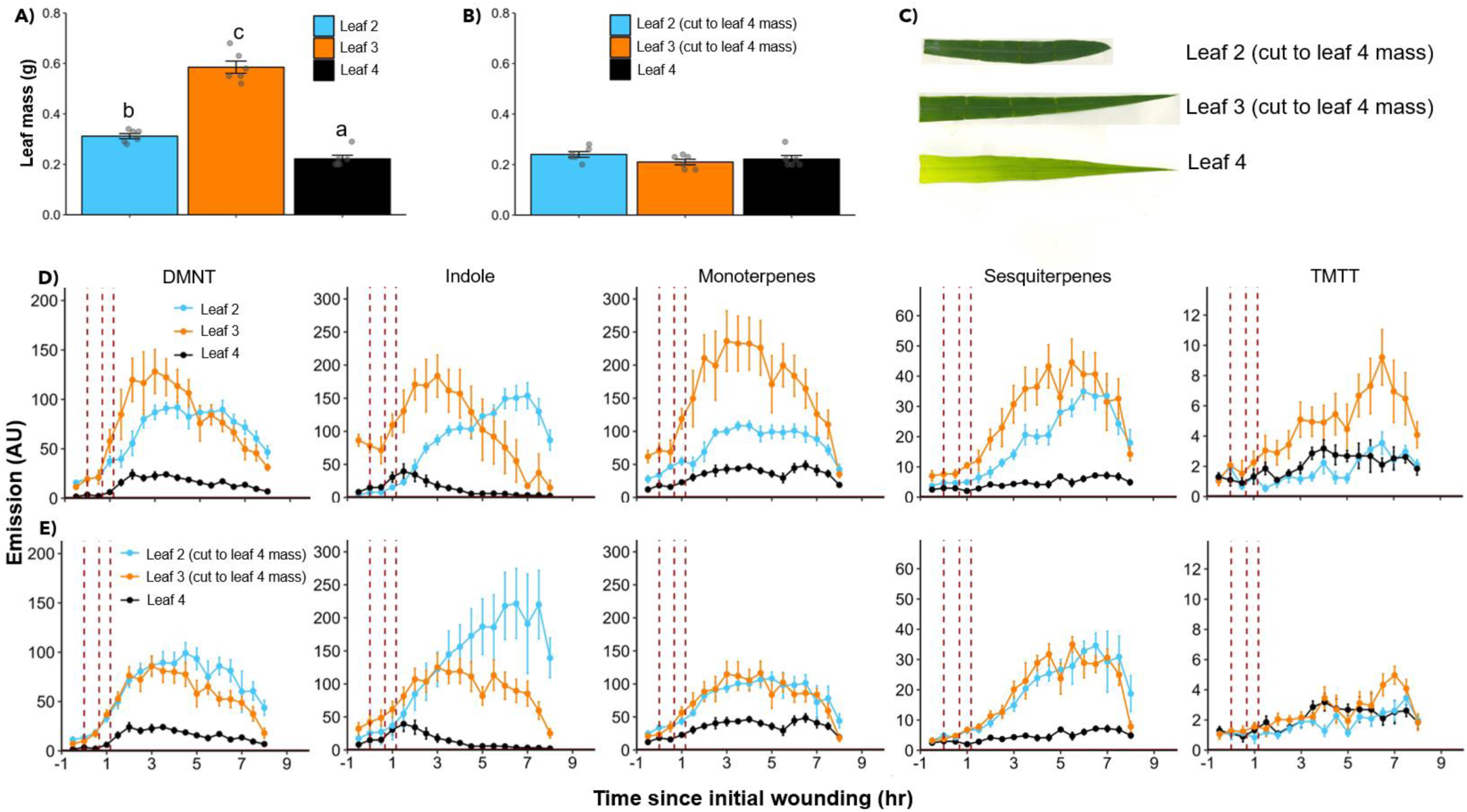
Standardizing leaf biomass results in similar induced volatile emissions between detached leaves of V2-stage seedlings. Biomass of A) entire detached leaves of V2-stage seedlings and B) V2-stage leaves 2 and 3 cut to the biomass of Leaf 4. C) V2-stage leaves 2-4 when standardized to size. Emission kinetics of D) entire detached V2-stage leaves and E) V2-stage leaves cut to leaf 4 biomass. Perforated vertical lines depict the timing of each mechanical damage entry. Lowercase letters indicate significant differences between leaves based on multiple comparisons tests. Abbreviations: DMNT, 4,8-dimethylnona-1,3,7-triene; TMTT, 4,8,12-trimethyltrideca-1,3,7,11-tetraene. Compounds were identified based on their molecular weight + 1, as all compounds were protonated. DMNT: m/z = 151.15, Indole: m/z = 118.07, Monoterpenes: m/z = 137.13, Sesquiterpenes: m/z = 205.20, TMTT: m/z = 219.21. Grey points represent biological replicates. Bars = mean ± SE. *n* = 6.

**Supplemental Figure 5.**
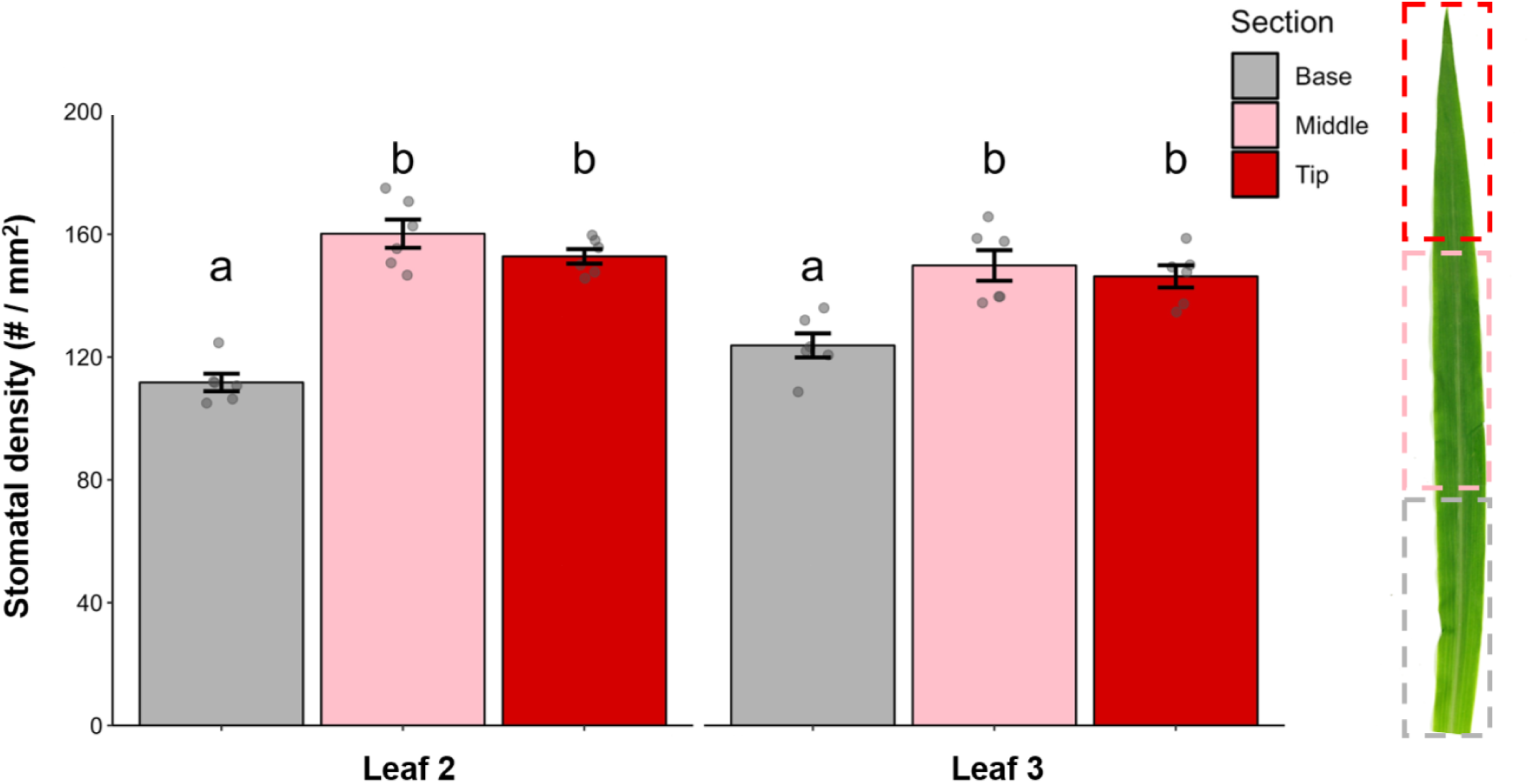
Stomatal patterns are similar between leaf 2 and leaf 3 in V2-stage maize seedlings. Stomatal density (abaxial and adaxial sides combined) of the sections of leaf 2 and leaf 3. Differing lowercase letters indicate significant differences between leaf and leaf section based on multiple comparisons tests. Grey points represent biological replicates. Bars = mean ± SE. *n* = 6.

### Generalist herbivore feeding patterns maximize volatile emissions

To test how herbivore feeding interacts with spatiotemporal differences in volatile emission, we placed a single *Spodoptera exigua* or *Spodoptera littoralis* larva on either leaf 2 or leaf 3 of V2-stage maize seedlings and tracked their feeding patterns. For *S. exigua*, we found that, regardless of starting position, herbivores consistently eat mostly leaf 3 (Fig 5). When placed on leaf 3, only 6.3% of plants had feeding on leaf 2 and 84.4% of plants had feeding damage on leaf 3 (Table 1). Even when placed on leaf 2, only 40.6% of plants had feeding damage on leaf 2 and 81.3% of plants had feeing on leaf 3. When herbivores were placed on leaf 2, damage area on leaf 4 was similar to leaf 2, and when placed on leaf 3, leaf 4 showed comparable damage levels to leaf 3. The trends for *S. littoralis* were similar to *S. exigua* with some key differences. Firstly, all *S. exigua* larvae placed on plants fed (*N* = 64). For *S. littoralis* only 36/64 fed (56%). *S. littoralis* also appeared far more mobile; many dropped off of into the soil at the base of the plant and did not eat. For *S. littoralis* larvae placed on leaf 2, although trends matched *S.exigua* patterns, differences in feeding area between leaves were not significant (Fig 5). In addition to damage area, for leaf 2-placed *S. littoralis*, the number of larvae feeding on each leaf was similar between leaves, whereas this was not the case for *S. exigua* placed on leaf 2 (Table 1). However, for larvae placed on leaf 3, the *S. littoralis* patterns matched those of *S. exigua*, whereby leaf 3 and leaf 4 had significantly more damage than the other leaves (Fig 5). Thus indicating that generalist herbivores preferentially feed on the leaves that result in the greatest plant-level damaged-induced volatile emissions.

**Figure 5.**
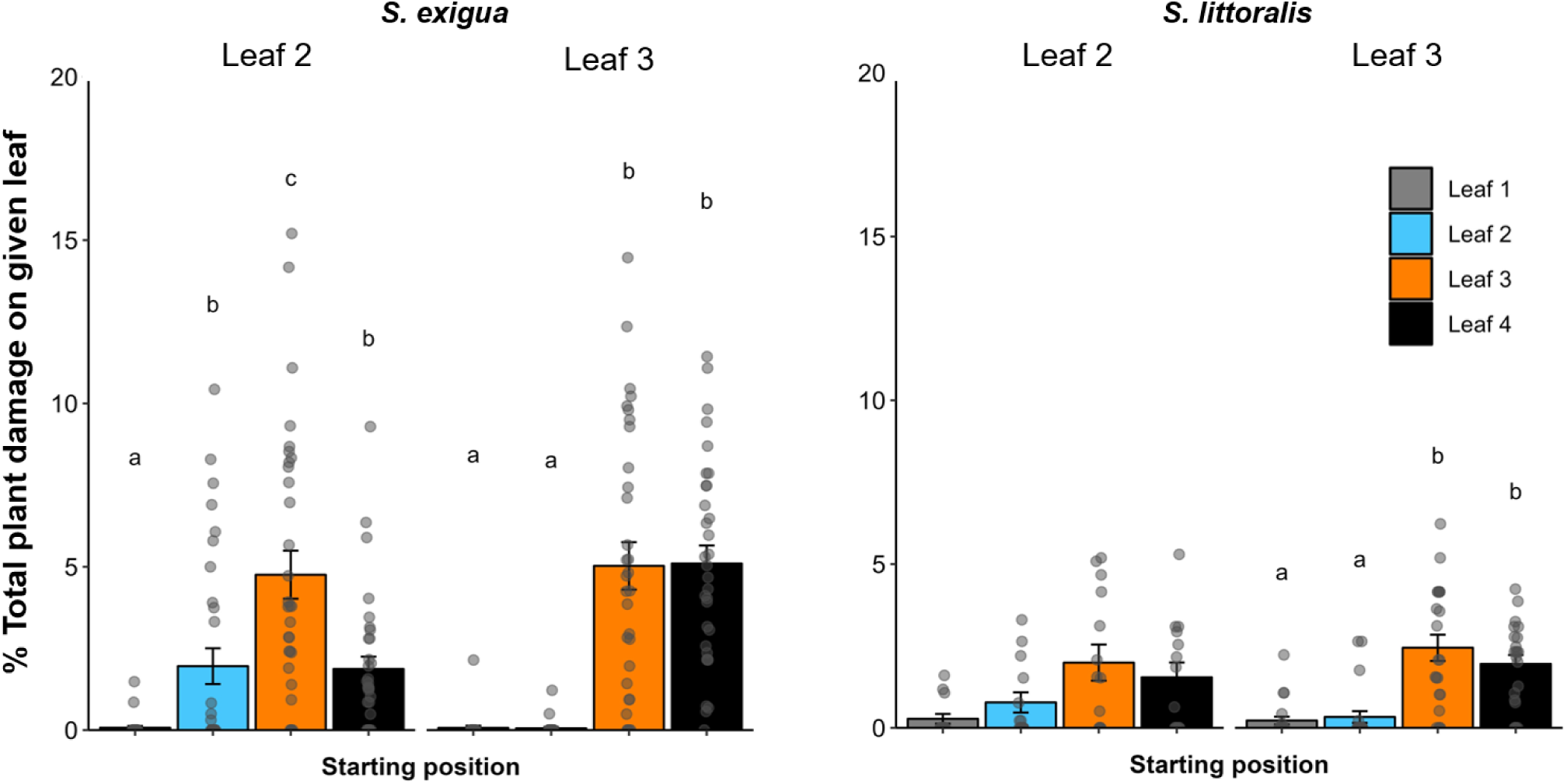
Generalist herbivores consistently feed heavily on leaf 3 regardless of starting position. The percentages of total plant damage (on V2-stage seedlings) done on each leaf for both *Spodoptera exigua* and *S. littoralis*. Larvae were either placed on leaf 2 or leaf 3 (starting position). For each herbivore species, lowercase letters indicate significant differences between leaves within a given herbivore starting position based on pairwise Wilcox tests. Grey points represent biological replicates. Bars = mean ± SE. For *S. exigua*, *n* = 32 and for *S. littoralis*, *n* = 14-22.

**Table 1.** Intact maize seedling herbivore feeding information table.

| Starting position | Leaf | # of larvae feeding (total) | % of larvae feeding |
| --- | --- | --- | --- |
| *Spodoptera exigua* |  |  |  |
| Leaf 2 | 1 | 2 (32) | 6.3 % |
|  | 2 | 13 (32) | 40.6 % |
|  | 3 | 26 (32) | 81.3 % |
|  | 4 | 23 (32) | 71.9 % |
| Leaf 3 | 1 | 1 (32) | 3.1 % |
|  | 2 | 2 (32) | 6.3 % |
|  | 3 | 27 (32) | 84.4 % |
|  | 4 | 31 (32) | 96.9 % |
| *Spodoptera littoralis* |  |  |  |
| Leaf 2 | 1 | 3 (14) | 21.4% |
|  | 2 | 7 (14) | 50 % |
|  | 3 | 9 (14) | 64.3 % |
|  | 4 | 8 (14) | 57.1 % |
| Leaf 3 | 1 | 5 (22) | 22.7 % |
|  | 2 | 4 (22) | 18.2 % |
|  | 3 | 18 (22) | 81.8 % |
|  | 4 | 18 (22) | 81.8 % |

Similar to mechanical damage patterns, we found, under authentic herbivory conditions, leaf 3 emitted higher amounts of all measured volatiles compared to leaf 2 in detached leaves (Fig 6A). In intact seedlings, total emissions from *S. exigua*-infested plants were positively correlated with leaf 3 damage for all measured compounds (Fig 6B): This was not the case for leaf 2, where no positive correlations were observed, and TMTT emission even showed a significant negative relationship with leaf 2 damage (Fig 6C). Thus confirming that, like mechanical damage, herbivory on leaf 3 is of the greatest consequence to total volatile emissions in maize seedlings.

**Figure 6.**
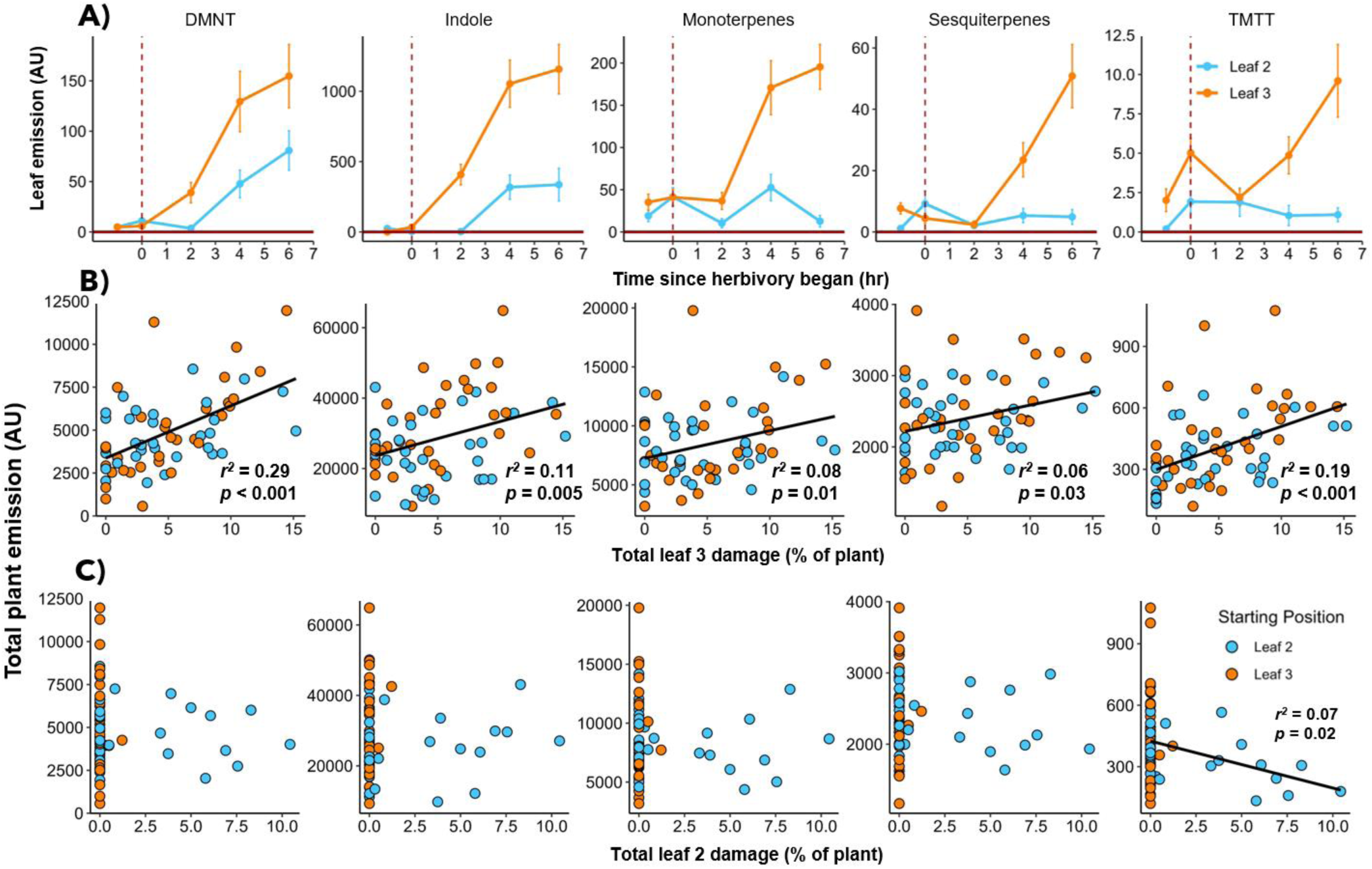
Leaf 3 plays a determinant role in overall herbivore-induced volatile emission during *Spodoptera exigua* herbivory. A) Emission from *S. exigua*-fed detached leaves from V2-stage seedlings. B) Correlation of total plant-level emission with total leaf 3 damage after 6 hr of herbivory. C) Correlation of total volatile emission with total leaf 2 damage after 6 hr of herbivory. B and C depict the same individual plants. For B and C, total emissions are the sum of individual measurement time points (taken every *ca.* 6 min) over 6 hr (see Materials and Methods, *Volatile sampling*). Colored points represent biological replicates and black lines represent significant linear relationships between emission and damage.

In addition to monitoring how herbivores mediate volatile emission patterns, we aimed to investigate how maize leaves differentially impact herbivore performance. Both *S. exigua* and *S. littoralis* damaged leaf 3 to a greater extent than leaf 2 in detached leaves (Fig 7A). After 48 hr of feeding, there were no clear differences in relative growth rate for either species between leaf 2- and leaf 3-fed larvae (Fig 7C).

**Figure 7.**
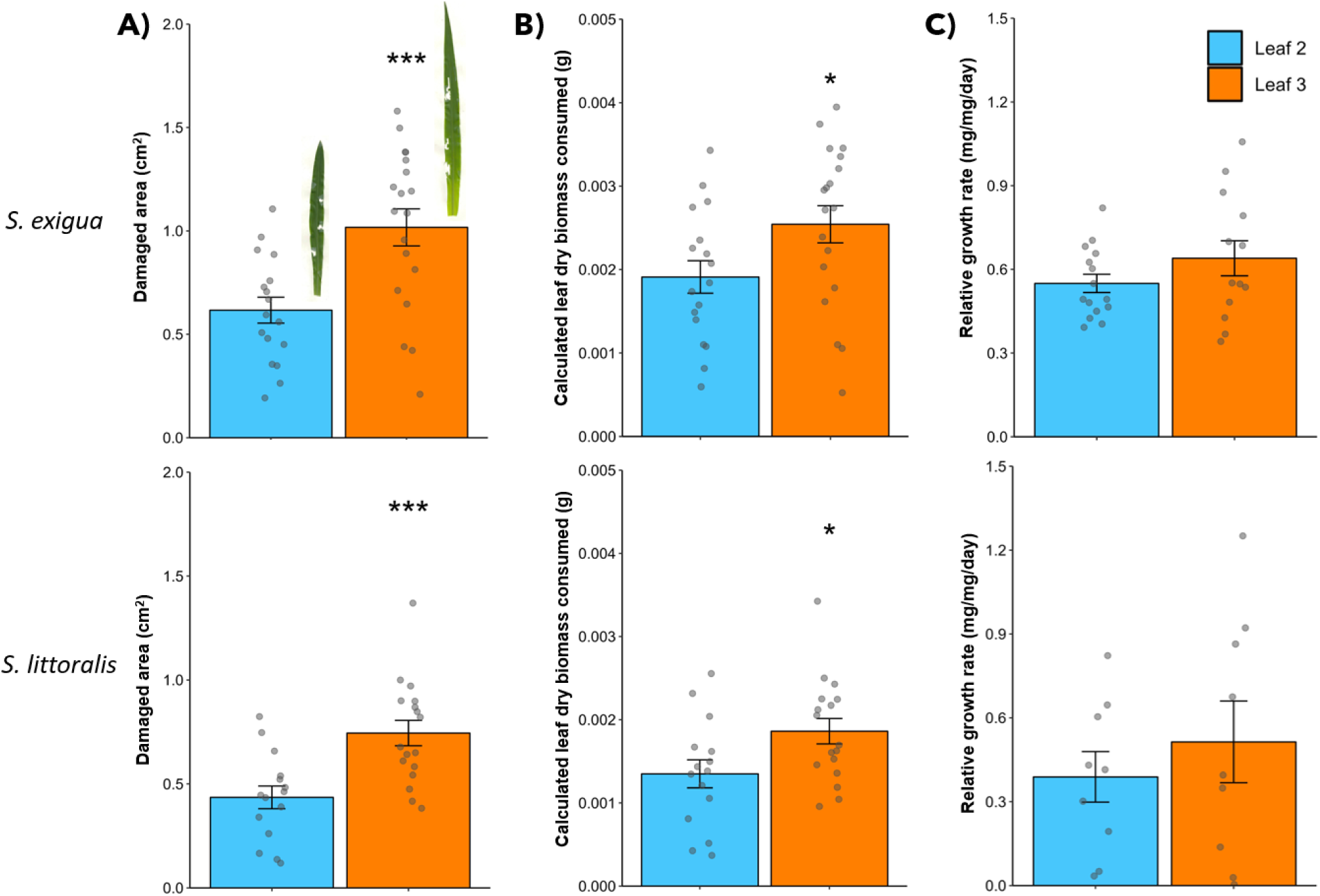
Generalist herbivores damage a greater area of leaf 3 with no apparent performance benefit. A) Area of leaf tissue consumed by 4th instar *Spodoptera exigua* and *S. littoralis* after 6 hr feeding on detached leaves from V2-stage seedlings, B) calculated biomass consumption based on leaf mass per area (LMA) of leaf 2 (0.0031) and leaf 3 (0.0025), and C) relative growth rate of 4th *S. exigua* and *S. littoralis* after 48 hr of feeding. * = *p* < 0.05, *** = *p* < 0.001 as determined by Welch’s two-sample *t-*test. Grey points represent biological replicates. Bars = mean ± SE. For *S. exigua n* = 13-19, for *S. littoralis* n = 9-17.

## Discussion

Spatiotemporal and developmental patterns have been shown influence volatile emissions in ecologically important ways (Köllner *et al*., 2013; Wang *et al*., 2023; Waterman *et al*., 2024). We show that size of the damaged maize leaves plays a determinant role in the overall plant-level volatile emissions and that generalist herbivores preferentially feed on the leaves that result in the highest volatile emissions. Here we discuss the mechanisms and potential implications of these patterns.

Within-plant spatiotemporal patterns are known to be important factors mediating stress responses (Barton *et al*., 2010; Köhler *et al*., 2015; Wang *et al*., 2023). Given maize seedlings differentially emit volatiles depending on the location of damage, one possible explanation for this variability could be differing induction capacities or systemic signaling capabilities between foliar tissues (Köllner *et al*., 2013; Irmisch *et al*., 2014). However, our data show minimal differences in induction between leaves; volatile biosynthesis genes are not only comparable across locally-damaged leaves, but the capacity for induction in undamaged, systemic leaves are consistent regardless of where damage occurs. A similar pattern was found for HDMBOA-Glc, which is a compound known to play an important role in direct defense against chewing herbivores such as *Spodoptera* spp. (Glauser *et al*., 2011; Tzin *et al*., 2017; Li *et al*., 2018). As such, per unit tissue, induction capacity is similar between V2-stage maize seedling leaves, most clearly shown between leaf 2 and leaf 3. Nevertheless, it is possible that underdeveloped defense machinery and gas exchange capacity in the emerging leaf 4 contributes to the apparent reductions in volatile emissions in leaf 4 (Wang *et al*., 2019; Wang *et al*., 2023). Based on these findings, it is clear that the observed differences in emission are not driven by differences in defense inducibility between leaves.

Leaf size seems to be the determinant factor controlling volatile emissions in maize seedlings, as there are very strong and consistent relationships between emission and both damaged-leaf mass and area across developmental stages. While there is generally a strong relationship between emission and area, it is less robust as the relationship between emission and biomass. As such, it is likely that emission is a function of both area, whereby greater area equates to more points of exit for volatiles (e.g., more stomata), and thickness, whereby thicker leaves may have more tissue capable of producing these volatiles. Biomass is a function of both area and thickness, which likely explains why this relationship is strongest.

The relationship between emission and leaf size is also clearly strongest for leaves that are actively developing. As developing leaves show greater variation in mass and area than fully-developed leaves (Powell & Lenhard, 2012), the close relationship between size and emission is apparent. Once leaves are fully developed there is much lower variation in size between leaves, however there are still differences in emission between individual leaves. We have previously identified leaf development as a factor that influences volatile emissions (Wang *et al*., 2023), and our findings in the present study support this; when immature and mature leaves are comparable in size, mature leaves emit more volatiles given equal damage.

While green leaf volatiles are formed in, and emitted directly from, wounded tissues (Hatanaka, 1993; Matsui *et al*., 2000; Escobar-Bravo *et al*., 2023), other volatiles such as terpenes and indole are produced *de novo* and systemically over time following stress (Erb *et al*., 2015), and emissions of these compounds are likely regulated by stomatal aperture (Niinemets, 2003; Seidl-Adams *et al*., 2015; Maleki *et al*., 2024). Some herbivores have been shown to excrete enzymes, such as glucose oxidase, which close stomata and minimize volatile emissions during feeding (Lin *et al*., 2021). Nevertheless, the relationship between herbivore feeding behavior and volatile emissions is not well understood. We find that two generalist herbivores preferentially feed on the leaves that result in the highest local and plant-level volatile emissions. As herbivores feed more heavily on leaf 3, which emits the most volatiles, we reasoned that there would be a clear benefit for herbivores to eat this leaf. However, surprisingly, this was not the case. In fact, what we observed was that, in order to reach the same level of growth, herbivores must consume more leaf 3 tissue compared to leaf 2, resulting in even greater volatile emissions. Even in intact plants, where herbivores can move freely over the entire plant, herbivores chose to consume leaf 3 most, which has a determinant impact on overall volatile emissions in maize seedlings. As a result, *Spodoptera* larvae may reveal themselves to the environment by feeding on the tissues that release the most volatiles. Herbivore-induced volatile emissions can be disadvantageous for herbivores through direct toxicity (Veyrat *et al*., 2016; Chen *et al*., 2021; Chen *et al*., 2023), enhanced attraction of natural enemies (Michereff *et al*., 2019) and population and community resistance by enhancing neighboring plant defenses (Escobar-Bravo *et al*., 2023; Waterman *et al*., 2024). On the other hand, they can also be advantageous for herbivores by confusing natural enemies and can alter competition between herbivores by enhancing host location and increasing oviposition (Poelman *et al*., 2012; Signoretti *et al*., 2012; Zu *et al*., 2020). Additionally, it has been shown that relative ratios and blends of emitted compounds, beyond absolute amounts can shape volatile-mediated interactions between plants and the environment (Allmann & Baldwin, 2010; Fouchier *et al*., 2018). More research, including longer-term feeding experiments under natural conditions, is required to understand to the consequences of increased volatile emissions in the atmosphere, as well as the potential fitness costs and benefits of herbivore feeding behavior in this context.

In conclusion, our work shows that the size of damaged leaves determines the extent of damage-induce volatiles emitted from maize seedlings. While tissue damage can be inflicted by many abiotic and biotic stressors, it is particularly noteworthy that, in addition to mechanical damage, patterns are consistent with real herbivore feeding. Our work provides a basis for developing a deeper understanding of the precise nature of herbivore feeding patterns and defense phenotypes, and how the interaction of the two can yield potentially important ecological outcomes for both plants and herbivores.

## Supplemental Tables

**Supplemental Table 1.**
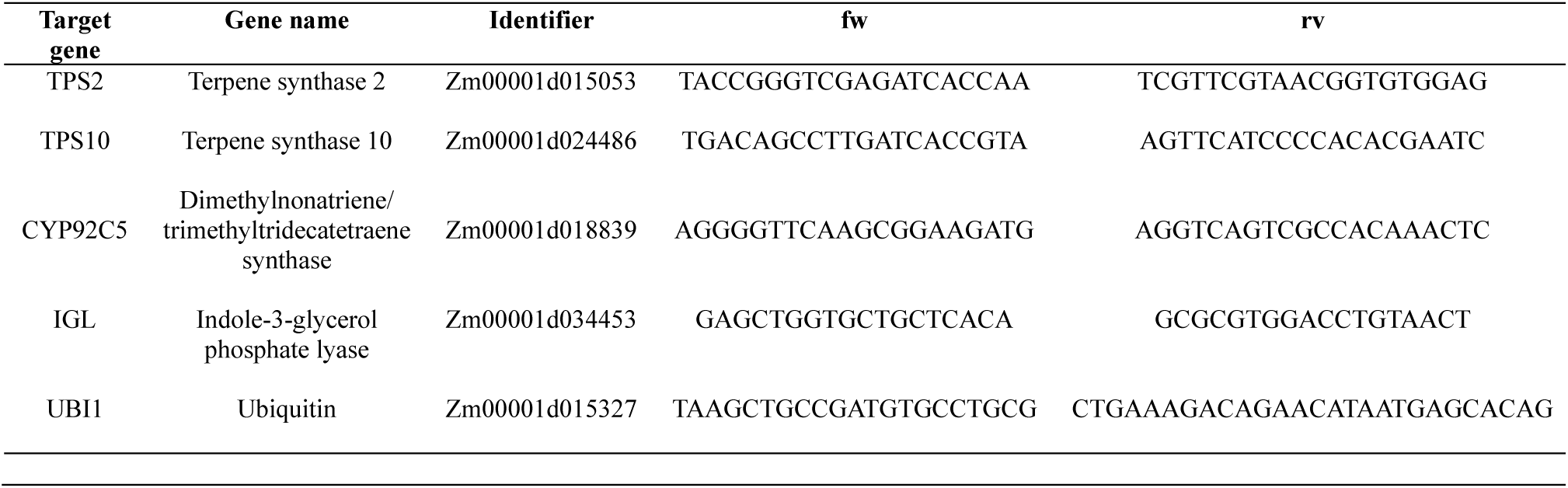
Gene identifiers and qRT-PCR primer sequences used in this study.

**Supplemental Table 2.** ANOVA and Kruskal-Wallis test results from data presented in the main text. Bold values: *p* < 0.05. Abbreviations: TPS2 = terpene synthase 2, TPS10 = terpene synthase 10, CYP92C5 = Dimethylnonatriene/trimethyltetradecatetraene synthase, IGL = indole-3-glycerol phosphate lyase, DMNT = 4,8-dimethylnona-1,3,7-triene, MNT = monoterpenes, SQT = sesquiterpenes, TMTT = 4,8,12-trimethyltrideca-1,3,7,11-tetraene, *S.e* = *Spodoptera exigua*, *S.l* = *Spodoptera littoralis*

|  | Damage location |  |  | Leaf |  |  | Dam x Leaf |  |  |
| --- | --- | --- | --- | --- | --- | --- | --- | --- | --- |
|  | *F* | *p* | *df* | *F/χ<sup>2</sup>* | *p* | *df* | *F* | *p* | *df* |
| *Gene expression*<br>(Fig 2) |  |  |  |  |  |  |  |  |  |
| TPS2 | 135.68 | < **0.001** | 3 | 10.65 | < **0.001** | 2 | 11.49 | < **0.001** | 6 |
| TPS10 | 454.67 | < **0.001** | 3 | 0.88 | 0.42 | 2 | 16.16 | < **0.001** | 6 |
| CYP92C5 | 87.16 | < **0.001** | 3 | 1.74 | 0.19 | 2 | 1.73 | 0.13 | 6 |
| IGL | 82.82 | < **0.001** | 3 | 6.64 | **0.003** | 2 | 2.55 | **0.03** | 6 |
| *Leaf mass*<br>(Fig 4A-B) |  |  |  |  |  |  |  |  |  |
| Entire leaves | - | - | - | 119.72 | < **0.001** | 2 | - | - | - |
| Cut leaves | - | - | - | 1.53 | 0.25 | 2 | - | - | - |
| *Plant damage*<br>(Fig 5) |  |  |  |  |  |  |  |  |  |
| *S.e* leaf 2 | - | - | - | 44.10 | < **0.001** | 3 | - | - | - |
| *S.e* leaf 3 | - | - | - | 89.39 | < **0.001** | 3 | - | - | - |
| *S.l* leaf 2 | - | - | - | 8.31 | **0.04** | 3 | - | - | - |
| *S.l* leaf 3 | - | - | - | 36.86 | < **0.001** | 3 | - | - | - |

**Supplemental Table 3.** Welch’s two-sample t-test results for herbivore performance parameters between detached leaf 2 and leaf 3.

| Response | *Spodoptera exigua* |  |  | *Spodoptera littoralis* |  |  |
| --- | --- | --- | --- | --- | --- | --- |
|  | *t* | *p* | df | *t* | *p* | df |
| Damage | -3.67 | <0.001 | 31 | -3.79 | <0.001 | 29 |
| Mass con. | -2.13 | 0.04 | 33 | -2.26 | 0.03 | 29 |
| RGR | -1.27 | 0.22 | 18 | -0.73 | 0.48 | 13 |

## Author Contributions

JMW conceived the ideas and designed experiments with critical inputs and support from ME. JMW, TMC, OMVL, PM and LW conducted experiments. JMW wrote the first draft of the manuscript and JMW and ME wrote the final version of the manuscript. All authors gave final approval of the manuscript. The authors have no conflicts of interest to declare.

## Acknowledgements

We would like to thank Sarah Holder for assistance with plant growth and all members of the Biotic Interactions and Chemical Ecology groups at the University of Bern for helpful discussions. This work was supported by the Swiss National Science Foundation (Grants Nr. 210651 and 200355), the State Secretariat for Education, Research, and Innovation SERI (Project CANWAS), as well as the University of Bern.

## Data availability statement

All data will be made available (either as supplemental material or on a data repository) upon publishing the final version, or beforehand upon request.

## References

Allmann S, Baldwin IT. 2010. Insects Betray Themselves in Nature to Predators by Rapid Isomerization of Green Leaf Volatiles. Science 329: 1075–1078.

Barton KE, Koricheva J, Associate Editor: Tia-Lynn Ashman, Editor: Donald L. DeAngelis. 2010. The Ontogeny of Plant Defense and Herbivory: Characterizing General Patterns Using Meta-Analysis. The American Naturalist 175: 481–493.

Bellec L, Seimandi-Corda G, Menacer K, Trabalon M, Ollivier J, Lunel C, Faure S, Cortesero A-M, Hervé M. 2022. Factors driving the within-plant patterns of resource exploitation in a herbivore. Functional Ecology 36: 1700–1712.

Block AK, Vaughan MM, Schmelz EA, Christensen SA. 2019. Biosynthesis and function of terpenoid defense compounds in maize (*Zea mays*). Planta 249: 21–30.

Bustos-Segura C, González-Salas R, Benrey B. 2022. Early damage enhances compensatory responses to herbivory in wild lima bean. Frontiers in Plant Science 13.

Chen C, Chen H, Huang S, Jiang T, Wang C, Tao Z, He C, Tang Q, Li P. 2021. Volatile DMNT directly protects plants against *Plutella xylostella* by disrupting the peritrophic matrix barrier in insect midgut. eLife 10.

Chen H, Chen C, Huang S, Zhao M, Wang T, Jiang T, Wang C, Tao Z, Zhang Y, Wang Y et al. 2023. Inactivation of RPX1 in Arabidopsis confers resistance to Plutella xylostella through the accumulation of the homoterpene DMNT. Plant, Cell & Environment 46: 946–961.

Eisenring M, Meissle M, Hagenbucher S, Naranjo SE, Wettstein F, Romeis J. 2017. Cotton Defense Induction Patterns Under Spatially, Temporally and Quantitatively Varying Herbivory Levels. Frontiers in Plant Science 8.

Erb M, Veyrat N, Robert CAM, Xu H, Frey M, Ton J, Turlings TCJ. 2015. Indole is an essential herbivore-induced volatile priming signal in maize. Nat. Commun. 6: 6273.

Escobar-Bravo R, Lin P-A, Waterman JM, Erb M. 2023. Dynamic environmental interactions shaped by vegetative plant volatiles. Nat. Prod. Rep. doi: 10.1039/D2NP00061J.

Fouchier A de, Sun X, Caballero-Vidal G, Travaillard S, Jacquin-Joly E, Montagné N. 2018. Behavioral Effect of Plant Volatiles Binding to Spodoptera littoralis Larval Odorant Receptors. Front. Behav. Neurosci. 12.

Frey M, Stettner C, Paré PW, Schmelz EA, Tumlinson JH, Gierl A. 2000. An herbivore elicitor activates the gene for indole emission in maize. Proceedings of the National Academy of Sciences 97: 14801–14806.

Glauser G, Marti G, Villard N, Doyen GA, Wolfender J-L, Turlings TC, Erb M. 2011. Induction and detoxification of maize 1,4-benzoxazin-3-ones by insect herbivores. The Plant Journal 68: 901–911.

Gonin M, Lopez-Hilfiker F, Hutterli M, Erb M, Pflander M. 2018. AUTOSAMPLER.

Groen SC, Jiang S, Murphy AM, Cunniffe NJ, Westwood JH, Davey MP, Bruce TJA, Caulfield JC, Furzer OJ, Reed A et al. 2016. Virus Infection of Plants Alters Pollinator Preference: A Payback for Susceptible Hosts? PLOS Pathogens 12: e1005790.

Hajdu C, Molnár BP, Waterman JM, Machado RAR, Radványi D, Fónagy A, Khan SA, Vassor T, Biet B, Erb M et al. 2024. Volatile-mediated oviposition preference for healthy over root-infested plants by the European corn borer. Plant, Cell & Environment n/a.

Hatanaka A. 1993. The biogeneration of green odour by green leaves. Phytochemistry 34: 1201–1218.

Heil M, Silva Bueno JC. 2007. Within-plant signaling by volatiles leads to induction and priming of an indirect plant defense in nature. Proceedings of the National Academy of Sciences 104: 5467–5472.

Himanen SJ, Nerg A-M, Holopainen JK. 2009. Degree of herbivore feeding damage as an important contributor to multitrophic plant-parasitoid signaling under climate change. Plant Signaling & Behavior 4: 249–251.

Hunziker P, Lambertz SK, Weber K, Crocoll C, Halkier BA, Schulz A. 2021. Herbivore feeding preference corroborates optimal defense theory for specialized metabolites within plants. Proceedings of the National Academy of Sciences 118: e2111977118.

Irmisch S, Jiang Y, Chen F, Gershenzon J, Köllner TG. 2014. Terpene synthases and their contribution to herbivore-induced volatile emission in western balsam poplar (Populus trichocarpa). BMC Plant Biology 14: 270.

Kattge J, Díaz S, Lavorel S, Prentice IC, Leadly P, Bönisch G, Garnier E., Westoby M, Reich PB, Wright IJ et al. 2011. TRY – a global database of plant traits. Global Change Biology 17: 2905–2935.

Köhler A, Maag D, Veyrat N, Glauser G, Wolfender J-L, Turlings TCJ, Erb M. 2015. Within-plant distribution of 1,4-benzoxazin-3-ones contributes to herbivore niche differentiation in maize. Plant, Cell & Environment 38: 1081–1093.

Köllner TG, Lenk C, Schnee C, Köpke S, Lindemann P, Gershenzon J, Degenhardt J. 2013. Localization of sesquiterpene formation and emission in maize leaves after herbivore damage. BMC Plant Biology 13: 15.

Lange ES de, Laplanche D, Guo H, Xu W, Vlimant M, Erb M, Ton J, Turlings TCJ. 2020. Spodoptera frugiperda Caterpillars Suppress Herbivore-Induced Volatile Emissions in Maize. Journal of Chemical Ecology 46: 344–360.

Li B, Förster C, Robert CAM, Züst T, Hu L, Machado RAR, Berset J-D, Handrick V, Knauer T, Hensel G et al. 2018. Convergent evolution of a metabolic switch between aphid and caterpillar resistance in cereals. Science Advances 4: eaat6797.

Lin P-A, Chen Y, Chaverra-Rodriguez D, Heu CC, Zainuddin NB, Sidhu JS, Peiffer M, Tan C-W, Helms A, Kim D et al. 2021. Silencing the alarm: an insect salivary enzyme closes plant stomata and inhibits volatile release. New Phytologist 230: 793–803.

Lin P-A, Peiffer M, Felton GW. 2020. Induction of defensive proteins in Solanaceae by salivary glucose oxidase of Helicoverpa zea caterpillars and consequences for larval performance. Arthropod-Plant Interactions 14: 317–325.

Maag D, Dalvit C, Thevenet D, Köhler A, Wouters FC, Vassão DG, Gershenzon J, Wolfender J-L, Turlings TCJ, Erb M et al. 2014. 3-β-D-Glucopyranosyl-6-methoxy-2-benzoxazolinone (MBOA-N-Glc) is an insect detoxification product of maize 1,4-benzoxazin-3-ones. Phytochem. 102: 97–105.

Maag D, Erb M, Bernal JS, Wolfender J-L, Turlings TCJ, Glauser G. 2015. Maize Domestication and Anti-Herbivore Defences: Leaf-Specific Dynamics during Early Ontogeny of Maize and Its Wild Ancestors. PloS one 10: e0135722.

Maag D, Köhler A, Robert CA, Frey M, Wolfender J-L, Turlings TC, Glauser G, Erb M. 2016. Highly localized and persistent induction of Bx1-dependent herbivore resistance factors in maize. The Plant Journal 88: 976–991.

Maleki FA, Seidl-Adams I, Felton GW, Kersch-Becker MF, Tumlinson JH. 2024. Stomata: gatekeepers of uptake and defense signaling by green leaf volatiles in maize. Journal of Experimental Botany: erae401.

Matsui K. 2006. Green leaf volatiles: hydroperoxide lyase pathway of oxylipin metabolism. Current Opinion in Plant Biology 9: 274–280.

Matsui K, Kurishita S, Hisamitsu A, Kajiwara T. 2000. A lipid-hydrolysing activity involved in hexenal formation. Biochemical Society Transactions 28: 857–860.

Maurya AK, Patel RC, Frost CJ. 2020. Acute toxicity of the plant volatile indole depends on herbivore specialization. Journal of Pest Science 93: 1107–1117.

Michereff MFF, Magalhães DM, Hassemer MJ, Laumann RA, Zhou J-J, Ribeiro, Paulo E. de A., Viana PA, Guimarães, Paulo E. de O., Schimmelpfeng PHC, Borges M et al. 2019. Variability in herbivore-induced defence signalling across different maize genotypes impacts significantly on natural enemy foraging behaviour. Journal of Pest Science 92: 723–736.

Mithöfer A, Wanner G, Boland W. 2005. Effects of feeding *Spodoptera littoralis* on lima bean leaves. II. Continuous mechanical wounding resembling insect feeding is sufficient to elicit herbivory-related volatile emission. Plant Physiol. 137: 1160–1168.

Niinemets Ü. 2003. Controls on the emission of plant volatiles through stomata: A sensitivity analysis. J. Geophys. Res. 108.

Ohnmeiss TE, Baldwin IT. 2000. OPTIMAL DEFENSE THEORY PREDICTS THE ONTOGENY OF AN INDUCED NICOTINE DEFENSE. Ecology 81: 1765–1783.

Orrock JL, Guiden PW, Pan VS, Karban R. 2022. Plant induced defenses that promote cannibalism reduce herbivory as effectively as highly pathogenic herbivore pathogens. Oecologia 199: 397–405.

Pan VS, Wetzel WC. 2024. Neutrality in plant–herbivore interactions. Proceedings of the Royal Society B: Biological Sciences 291: 20232687.

Paul-Victor C, Züst T, Rees M, Kliebenstein DJ, Turnbull LA. 2010. A new method for measuring relative growth rate can uncover the costs of defensive compounds in Arabidopsis thaliana. New Phytologist 187: 1102–1111.

Poelman EH, Bruinsma M, Zhu F, Weldegergis BT, Boursault AE, Jongema Y, van Loon JJA, Vet LEM, Harvey JA, Dicke M. 2012. Hyperparasitoids Use Herbivore-Induced Plant Volatiles to Locate Their Parasitoid Host. PLOS Biology 10: e1001435.

Powell AE, Lenhard M. 2012. Control of Organ Size in Plants. Current Biology 22: R360–R367.

R Core Team. 2022. R: A language and environment for statistical computing. Vienna, Austria: R Foundation for Statistical Computing.

Raissig MT, Matos JL, Anleu Gil MX, Kornfeld A, Bettadapur A, Abrash E, Allison HR, Badgley G, Vogel JP, Berry JA et al. 2017. Mobile MUTE specifies subsidiary cells to build physiologically improved grass stomata. Science 355: 1215–1218.

Richter A, Schaff C, Zhang Z, Lipka AE, Tian F, Köllner TG, Schnee C, Preiß S, Irmisch S, Jander G et al. 2016. Characterization of Biosynthetic Pathways for the Production of the Volatile Homoterpenes DMNT and TMTT in *Zea mays*. Plant Cell 28: 2651–2665.

Rodriguez-Saona CR, Rodriguez-Saona LE, Frost CJ. 2009. Herbivore-Induced Volatiles in the Perennial Shrub, Vaccinium corymbosum, and Their Role in Inter-branch Signaling. Journal of Chemical Ecology 35: 163–175.

Schmelz EA, Alborn HT, Banchio E, Tumlinson JH. 2003. Quantitative relationships between induced jasmonic acid levels and volatile emission in Zea mays during Spodoptera exigua herbivory. Planta 216: 665–673.

Schnee C, Köllner TG, Held M, Turlings TCJ, Gershenzon J, Degenhardt J. 2006. The products of a single maize sesquiterpene synthase form a volatile defense signal that attracts natural enemies of maize herbivores. Proceedings of the National Academy of Sciences 103: 1129–1134.

Seidl-Adams I, Richter A, Boomer KB, Yoshinaga N, Degenhardt J, Tumlinson JH. 2015. Emission of herbivore elicitor-induced sesquiterpenes is regulated by stomatal aperture in maize (*Zea mays*) seedlings. Plant Cell Environ. 38: 23–34.

Signoretti AGC, Peñaflor, M. F. G. V., Bento JMS. 2012. Fall Armyworm, Spodoptera frugiperda (J.E. Smith) (Lepidoptera: Noctuidae), Female Moths Respond to Herbivore-Induced Corn Volatiles. Neotropical Entomology 41: 22–26.

The Herbivory Variability Network* †, Robinson ML, Hahn PG, Inouye BD, Underwood N, Whitehead SR, Abbott KC, Bruna EM, Cacho NI, Dyer LA et al. 2023. Plant size, latitude, and phylogeny explain within-population variability in herbivory. Science 382: 679–683.

Turlings TC, Erb M. 2018. Tritrophic Interactions Mediated by Herbivore-Induced Plant Volatiles: Mechanisms, Ecological Relevance, and Application Potential. Annual Review of Entomology 63: 433–452.

Tzin V, Hojo Y, Strickler SR, Bartsch LJ, Archer CM, Ahern KR, Zhou S, Christensen SA, Galis I, Mueller LA et al. 2017. Rapid defense responses in maize leaves induced by Spodoptera exigua caterpillar feeding. Journal of Experimental Botany 68: 4709–4723.

Underwood N. 2012. When herbivores come back: effects of repeated damage on induced resistance. Functional Ecology 26: 1441–1449.

van den Berg J, Britz C, Du Plessis H. 2021. Maize Yield Response to Chemical Control of Spodoptera frugiperda at Different Plant Growth Stages in South Africa. Agriculture 11.

Veyrat N, Robert CAM, Turlings TCJ, Erb M. 2016. Herbivore intoxication as a potential primary function of an inducible volatile plant signal. Journal of Ecology 104: 591–600.

Wang H, Guo S, Qiao X, Guo J, Li Z, Zhou Y, Bai S, Gao Z, Wang D, Wang P et al. 2019. BZU2/ZmMUTE controls symmetrical division of guard mother cell and specifies neighbor cell fate in maize. PLOS Genetics 15: e1008377.

Wang L, Jäggi S, Cofer TM, Waterman JM, Walthert M, Glauser G, Erb M. 2023. Immature leaves are the dominant volatile-sensing organs of maize. Curr. Biol. 33: 3679–3689.e3.

Waterman JM, Cazzonelli CI, Hartley SE, Johnson SN. 2019. Simulated Herbivory: The Key to Disentangling Plant Defence Responses. Trends Ecol. Evol. 34: 447–458.

Waterman JM, Cibils-Stewart X, Cazzonelli CI, Hartley SE, Johnson SN. 2021. Short-term exposure to silicon rapidly enhances plant resistance to herbivory. Ecology 102: e03438.

Waterman JM, Cofer TM, Wang L, Glauser G, Erb M, Rasmann S, Weigel D. 2024. High-resolution kinetics of herbivore-induced plant volatile transfer reveal clocked response patterns in neighboring plants. eLife 12: RP89855.

White H. 1980. A Heteroskedasticity-Consistent Covariance Matrix Estimator and a Direct Test for Heteroskedasticity. Econometrica 48: 817–838.

Zu P, Boege K, Del-Val E, Schuman MC, Stevenson PC, Zaldivar-Riverón A, Saavedra S. 2020. Information arms race explains plant-herbivore chemical communication in ecological communities. Science 368: 1377–1381.

